# Unlocking Genetic Diversity in Colombian Cassava Landraces for Accelerated Breeding

**DOI:** 10.1101/2025.06.30.662420

**Authors:** Kehan Zhao, Evan Long, Francisco Sanchez, Paul Chavarriaga, Grey Monroe

## Abstract

Cassava (*Manihot esculenta* Crantz) is a staple food for hundreds of millions across the global south. In this study, we investigated genomic diversity among over 1000 cassava genotypes, with a particular focus on the addition of 387 newly sequenced landrace varieties originating from diverse climates across Colombia. As cassava was domesticated in or near the Amazon basin, these landraces represent untapped genetic diversity that could be used to help improve modern varieties. As theory predicts, we found that landraces retain high genetic diversity, observing variation lacking in breeding lines from Asia and Africa, where introductions likely caused population bottlenecks. Genetic differentiation in landraces reflects both space and climate of origin, suggesting the combined effects of demography and selection. To identify alleles with the potential to inform targets for gene editing, we assessed the diversity of loss-of-function (LoF) mutations across these landraces. We found evidence that deleterious LoF alleles were purged by inbreeding. Notably, genes retaining LoF alleles despite inbreeding were significantly enriched for functions related to the biosynthesis of coumarins and the regulation of plant immunity, suggesting selection on postharvest quality and disease resistance. We further identified specific loci associated with climates of origin, motivating future experiments using targeted knockouts to test hypotheses about the adaptive value of specific LoF alleles. This work supports longstanding hypotheses about landraces as a reservoir of genetic diversity and establishes the foundation to leverage this variation in cassava to discover alleles for accelerated breeding via gene editing.

**Short summary:** **This study explores the genetic diversity of cassava by sequencing 387 landrace varieties and wild relatives from diverse climates in Colombia, aiming to identify potential gene targets for gene editing to enhance climate resilience. The research focuses on loss-of-function mutations, which are expected to have large effects and provide testable targets. Genome-wide association analysis reveals multiple potential targets associated with climate adaptation in cassava**.

## Introduction

Cassava (*Manihot esculenta* Crantz), a versatile and resilient monoecious root crop, provides a major caloric source for over 500 million people worldwide, especially in the global south (Parmar *et al*., 2017; Ferguson *et al*., 2019). The importance of cassava in global agriculture is largely due to its remarkable productivity with minimal inputs and ability to thrive under marginal conditions, such as in water limited environments (Parmar *et al*., 2017). However, this adaptability faces new environmental challenges including extreme weather events and plant diseases (Food and Agriculture Organization, 2019). While cassava is a staple in regions of the world with the fastest human population growth, recent projections have predicted a possible drop in cassava yield of up to 10-20% in the next 50-100 years (Zhu *et al*., 2023). These underscore the necessity to fortify the resilience of essential crops like cassava for future food security.

Cassava was domesticated in or near the Amazon Basin between 5 and 10 thousand years ago, and has since become a major crop in the tropical regions of South America, Africa, and Asia (Parmar *et al*., 2017; Ferguson *et al*., 2019). In recent years, wild germplasm and domesticated accessions not used for intensive breeding have been seen as a valuable resource to access adaptive genetic diversity (Kashyap *et al*., 2022). Crop landraces, or traditional cultivated varieties, can be valuable sources of beneficial traits and alleles, often due to adaptation reflecting geographical variation. Leveraging landraces or wild relatives to identify genetic variation responsible for environmental adaptation has been successful in multiple crops and other plant species including wheat (Lopes *et al*., 2015), maize (Janzen *et al*., 2022), *Brassicaceae* (*Turner et al., 2010*), and sorghum (Lasky *et al*., 2015).

Here, we characterized the genomic diversity of Colombian cassava landraces maintained in the living genbank collection at the Alliance Bioversity and the International Center for Tropical Agriculture (CIAT). This genebank contains thousands of cassava clones, many of which were sampled and preserved from locations in Colombia and the rest of South America (Ferguson *et al*., 2019). These accessions originated from locations that represent diverse environments, especially across the dramatic elevation gradient of Colombia, and thus may also contain alleles responsible for local climate adaptation. As a first look at this genomic variation, we aim to provide foundational resources to discover loci informative for accelerated breeding.

The advent of genome engineering with technologies such as CRISPR-Cas9 presents an opportunity to match the pace of crop improvement with that of the challenges imposed by environmental change (Sedeek *et al*., 2019). This technology has already proved useful in cassava by inducing resistance to cassava brown streak disease, without relying on transgenics (Gomez *et al*., 2019). While mapping studies can find markers for certain traits, it remains difficult to reliably find causative mutations that can be the target of genome engineering (Haque *et al*., 2018). To identify actionable targets for gene editing, it is essential to consider the functional characterization of causative mutations. Loss-of-function (LoF) mutations, or gene knockouts, are classified as mutations predicted to disrupt or remove the function of a protein. These include premature stop codons, reading frame shifts, large deletions, or entire gene loss. Naturally occurring LoF mutations are important in the evolution of many plants, contributing to domestication, crop improvement, and adaptation to the environment (Orr, 2005; Monroe *et al*., 2016, 2018, 2020; Murray, 2020; Xu & Guo, 2020; Lee *et al*., 2024; Klim *et al*., 2024). Identifying beneficial LoF alleles in crops can thus inform next-generation breeding efforts for rapid improvement. Indeed, gene knockouts have already led to major agricultural advances. The Green Revolution’s dramatic yield increases were driven by LoF mutations in *GA20ox2* in rice and other crops (Spielmeyer *et al*., 2002). In wheat, knockout of *GRAIN WIDTH2* improves rust resistance and grain weight (Sestili *et al*., 2019; Liu *et al*., 2024), while in cassava, knocking out *CYP79D1* and *CYP79D2*, which are key genes in cyanide synthesis, has reduced cyanide levels by 92% in bitter varieties (Jørgensen *et al*., 2005).

However, detecting causal gene loss using statistical approaches such as Genome-Wide Association Study (GWAS) can be challenging as allelic heterogeneity diminishes the power of statistical testing (Monroe *et al*., 2021). To address this, LoF burden tests aggregate multiple rare or independent LoF variants within a gene into a single score, enhancing the ability to detect associations between gene disruption and phenotypic variation (Povysil *et al*., 2019; Spence *et al*., 2024). In this study, we sequenced and analyzed 387 cassava landrace genomes, together with previous whole genome sequencing of wild relatives and improved breeding lines. We tested for genetic-enviornment associations at multiple levels, including loss-of-function alleles. This approach identifies multiple candidate genes linked to climate adaptation and offers new directions for breeding strategies in cassava.

## Materials and Methods

### Landrace Sampling

We selected 387 unique Cassava landraces based on their environmental parameters across Colombia to best represent the geological diversity within the country. We also sequenced an additional 33 technical duplicates of some of the chosen accessions to verify and ensure sequencing and variant calling accuracy. Tissue cultures of those selected accessions were sent to the laboratory for DNA extraction from the CIAT genebank.

### DNA Isolation and Sequencing

Young leaf tissue from each accession was sampled to extract genomic DNA using the DNeasy plant mini kit (QIAGEN). DNA samples were sequenced with DNBSEQ™ at BGI America, yielding a total of ∼2.7 × 10^10^ 150-bp paired-end reads and an average depth of coverage at 25.20× for each sample.

### Variant Calling

We downloaded previously published Cassava whole-genome sequencing data from Ramu *et al*. (2017), Kistler *et al*. (2025) (excluding herbarium and archaeological samples), Hu *et al*. (2021), Bredeson *et al*. (2016), and Wang *et al*. (2014), and conducted *de novo* variant calling along with our sequencing data of Colombian cassava landraces and close relatives. Raw reads were trimmed using Trimmomatic (version 0.39) to control read quality. The clean reads were mapped to the reference genome v8 (https://phytozome-next.jgi.doe.gov/info/Mesculenta_v8_1) using BWA-MEM (version 0.7.17-r1188). The mapped reads were then sorted and duplicates were removed by samtools (version 1.13). The reads were realigned and the variants were called for each accession using DeepVariant (version 1.5.0, Poplin *et al*., 2018). Subsequently, gVCF merging and joint variant calling was performed using GLnexus (version 1.4.1, Yun *et al*., 2021). To determine the effects of variants on the genome, we used SnpEff (version 5.1d, Cingolani *et al*., 2012) to build a database from the reference genome (v8) and annotated the VCF.

### Data Filtering and Quality Control

We employed a combination of novel and standard filtering methods to obtain high-confidence variants. Among our genotyped individuals we obtained 33 technical replicates (i.e. 33 individuals whose DNA was sampled, extracted, and sequenced in duplicate). We used the sites with discordance between the two replicate genotype calls as a training set to train a gradient boost model (R package XGboost, Chen & Guestrin, 2016) to detect erroneous variant sites. In brief, a set of variant site characteristics (percent missing genotypes, Quality score, minor allele frequency, number of alleles, and depth mean and variance) to predict erroneous sites. The percent missing genotypes was the most significant predictor of discordant sites between technical replicates. This was done in a Leave-one-out method for each chromosome and the predicted erroneous sites were removed from the available variants. The variants were further filtered using more standard methods using vcftools (version 0.1.16), sites with Quality value above 30 (−-minQ 30), Genotype Quality value above 20 (−-minGQ 20), and Minor Allele Frequency above 0.05 (−-maf 0.05) were kept for downstream analysis. Variants were then phased using BEAGLE V5 (Browning & Browning, 2007) with the ne=1000.

### Assessing Genetic Diversity and Population Structure

Principal Component Analysis (PCA) was performed on the filtered VCF using Plink (version 1.9). Genetic assignment analysis was conducted using the filtered SNPs with the ADMIXTURE program (Version 1.3.0, Alexander *et al*., 2009). The sequencing data from Hu *et al*. (2021) had an average depth of ∼8.39×, while our newly sequenced Colombian landraces were generated at a depth of 25×. Accessions from Kistler *et al*. (2025) had an average depth of 21.31×, but ranged widely from 0.01× to 88.01×. These substantial differences in sequencing depth may significantly affect variant calling, data filtering, and downstream analyses. Therefore, accessions from Hu *et al*. (2021) and Kistler *et al*. (2025) were excluded from subsequent analyses of linkage disequilibrium (LD), nucleotide diversity (π), and fixation index (F_ST_) comparisons.

To estimate nucleotide diversity unbiased by missing data in the VCF, all-site VCFs (VCF including invariant sites) of 18 chromosomes were produced using GATK HaplotypeCaller, GenomicsDBImport, and GenotypeGVCFs (version 4.5.0.0). F_ST_ and π between those three groups were then calculated using Pixy (1.2.10.beta2, Korunes & Samuk, 2021) after basic filtering (−-max-missing 0.8 \ --min-meanDP 20 \ --max-meanDP 500). Linkage disequilibrium was also calculated using Plink (version 1.9) on a window size of 100 kb.

### Environmental Variables and Phenotypes

Environmental variables were collected using the latitude and longitude of the sampled cassava landraces. Bioclim variables (Fick & Hijmans, 2017) were extracted using the R package “raster” version 3.6.26, with a tile resolution of 2.5. Elevation values were extracted using the R package “elevatr” version 0.99.0. Composite environmental phenotypes were generated using principal components analysis on all environmental variables corresponding to EnvPC1-5. Phenotypic data were previously collected and reported by the CIAT genebank, provided as metadata for each accession.

### Annotating Loss-of-Function Mutations

We used mutation effect and protein structure prediction to evaluate loss-of-function (LoF) mutations and mutation impact prediction to infer potential loss-of-function mutations. We used the primary transcripts annotated from the version 8 cassava genome assembly (Nordberg *et al*., 2014) for all gene analyses. The program SnpEff (Cingolani *et al*., 2012) was used to annotate mutation effects from our genotype variants. All mutations classified as “HIGH” effect were putative LoF mutations, including mutations such as frameshift insertions and deleterious, gain of a premature stop codon, loss of the start codon, and splice interruption.

To additionally validate the impact of LoF mutations on protein structure, we performed protein structure prediction using ESM-fold (Lin *et al*., 2023). We used two values of amino acid structure prediction to measure their characteristics, the confidence or “predicted local distance difference test” score (pLDDT) and the relative available surface area (rASA). These protein characteristics were measured across all proteins using a github pipeline https://github.com/em255/PopulationPDBStats. These protein structures often have 5’ and 3’ tails that are highly disordered and predicted with low confidence, where we see an enrichment for mutations. We annotated the 5’ tail of a protein as the region starting from the first amino acid until the first confidently folded amino acid occurs within an ordered region of the protein (pLDDT >70 & rASA<0.5). The 3’ tail was annotated in a similar manner, starting from the 3’ end. Using these disordered tail annotations, we chose to disregard certain LoF mutations that we estimated not to impact overall protein structure. All mutations in the disordered 3’ tail were ignored. Premature stop mutations and start codon losses were disregarded in the 5’ tail if another in-frame start codon was present within the region. While this method may discount some functional importance of these disordered regions, it allows for the enrichment of likely high-impact LoF mutations. Finally, using phased genotype information multiple LoF mutations were collapsed into a single functional status for each gene with either 0, 1 (Heterozygous), or 2 functional copies of the gene. Importantly, rare alleles (MAF < 0.05) were included into this functional genotyping.

### Genome-Wide Association

Genome-wide association was performed using tassel-5 (Bradbury *et al*., 2007) utilizing either all variants or LoF matrix as genotype predictors. Each phenotype was fit using the generalized linear model function of tassel with the first five genetic principal components as fixed covariates to control for population structure. For association using LoF status, a minor LoF frequency of 5% was imposed similar to minor allele frequency.

To further test for the effect of including population structure on association detection, we conducted association analyses in R using linear models which included as covariates the principal components of genetic kinship among genotypes. From these results, we then identified the genes showing significant associations between functional status and environmental variables after false discovery rate correction of p-values.

### Redundancy Analysis

We performed redundancy (RDA) and partial redundancy analysis (pRDA) following published tutorials (Capblancq & Forester, 2021). We applied this method to our genotype and environmental data to evaluate the genetic variance explained by environmental predictors. Environmental predictors were standardized to ensure comparability and then used to estimate the total variance explained of the genetic relationships. We then used pRDA to partition the variance explained by climate, neutral genetic structure, and geography. This involved creating models with population allele frequencies as the response variable and sets of bioclimate variables, genetic structure proxies, and geographical coordinates as explanatory variables. This approach allowed us to decompose the contributions of different factors to genetic variation and assess their independent and combined effects.

Loci with p-values below a stringent threshold were identified as candidate adaptive outliers. Results were visualized using an RDA biplot and a Manhattan plot, showing the correlation between genetic variation and environmental predictors, and highlighting significant outliers. This approach allowed us to identify loci potentially under selection due to environmental factors while accounting for population structure, providing insights into the genetic basis of local adaptation.

### Gene Ontology

To investigate the biological significance of LoF mutations, we identified putative orthologs of cassava (*Manihot esculenta*) genes in *Arabidopsis thaliana* (TAIR10) using BLASTP, retaining matches with an E-value <1e-10 and bit score >100. For each cassava gene containing at least one predicted LoF allele, we mapped its Arabidopsis ortholog and used the associated Gene Ontology (GO) annotations to perform enrichment analysis, focusing on the Biological Process (BP) category. Enrichment was conducted using the TopGO package in R, applying the “weight01” algorithm and Fisher’s exact test. Only GO terms that remained significant after Bonferroni correction for multiple testing (adjusted p < 0.05) were considered enriched. To assess lineage-specific patterns, we performed the analysis separately for wild relatives, traditional landraces, and modern breeding lines.

### Testing for Selection and Inbreeding Effects on Loss-of-Function Accumulation

To evaluate whether deleterious LoF mutations are purged by negative selection under inbreeding, we tested for an association between LoF burden and genomic inbreeding coefficients in cassava landraces. For each individual landrace, we quantified the total number of LoF events, defined as the number of genes with at least one high-confidence LoF allele. In parallel, we estimated the inbreeding coefficient (F_IS_) using VCFtools, based on genome-wide biallelic SNP variation. We then assessed the relationship between inbreeding and LoF accumulation by ranking landraces according to their F_IS_ values and their total number of LoF genes. A negative correlation between ranked F_IS_ and LoF counts would support the hypothesis that inbreeding facilitates purging of deleterious variants through increased homozygosity and stronger exposure to selection.

To investigate whether certain LoF mutations may be beneficial in specific functional contexts, we repeated this analysis while restricting LoF counts to genes annotated with specific Gene Ontology (GO) Biological Process categories. This allowed us to identify cases where LoF accumulation is positively associated with inbreeding, which may indicate adaptive or tolerated LoF in particular functional pathways.

## Results

### Genetic Diversity and Population Structure

To investigate the genetic variation of Cassava landraces in Colombia, we sampled and sequenced 387 unique cassava accessions by whole-genome sequencing (WGS). On average, 25.20x coverage was generated for each accession. We then combined our data with previously published cassava WGS data, including 174 accessions from Ramu *et al*. (2017), 158 accessions from Kistler *et al*. (2025), 338 accessions from Hu *et al*. (2021), 59 accessions from Bredeson *et al*. (2016) and 2 accessions from Wang *et al*. (2014), and conducted *de novo* variant calling (**Table S1**). To ensure consistency and minimize batch effects, all samples were processed using the same quality control, filtering, and variant calling pipeline (***Materials and methods***).

To dissect the population structure underlying cassava samples, we performed principal component analysis (PCA) of selected high-confidence variants, which reveals substantial, previously untapped genetic diversity among Colombian cassava landraces (**Fig. 1**). Within *Manihot esculenta*, a total of 29.23% genetic variance was captured in the first two principal components (PCs). South American accessions displayed greater genetic differentiation compared to cultivated cassava from Africa and Asia, although some overlap was observed, suggesting a genetic bottleneck associated with the introduction of cassava to these regions (**Fig. 1b**). Notably, PC1 showed a strong negative correlation with elevation (**Fig. 2a, Fig. S1**) and a positive correlation with temperature (**Fig. 2b**), indicating that the genetic differentiation along PC1 may reflect local adaptation or environmental selection associated with altitude and climate. Highland and lowland populations might harbor distinct allelic compositions, possibly tied to important traits such as drought and heat tolerance along with demographic history.

**Figure 1.**
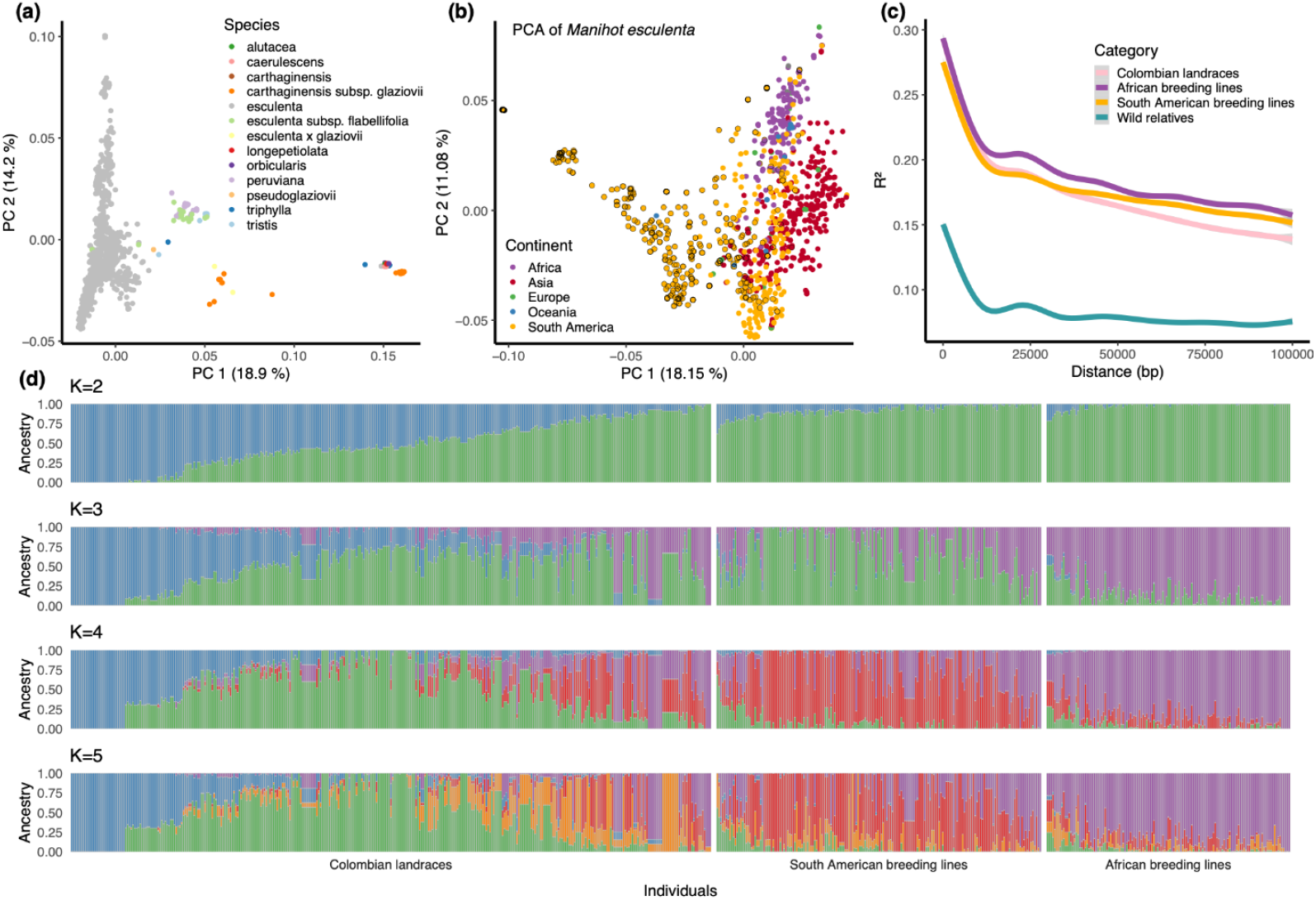
Population Structure, Genetic Diversity, and Linkage Disequilibrium Decay of Cassava Accessions and Close Relatives. **(a)** Principal component analysis (PCA) of the first two dimensions of genotype data from all accessions, including close relatives. Data points are colored by species. **(b)** PCA of the first two dimensions of genotype data from *Manihot esculenta* accessions, with data points colored by geographical location (continent). Colombian landrace accessions sequenced in this study are highlighted in black outline. **(c)** Linkage disequilibrium (LD) decay in Colombian landraces, African breeding lines, South American breeding lines, and wild relatives. **(d)** ADMIXTURE plots of Colombian landraces, African breeding lines, and South American breeding lines. Cassava accessions progressively separate into minor subpopulations as K increases. Accessions are arranged in the order of PC1.

**Figure 2.**
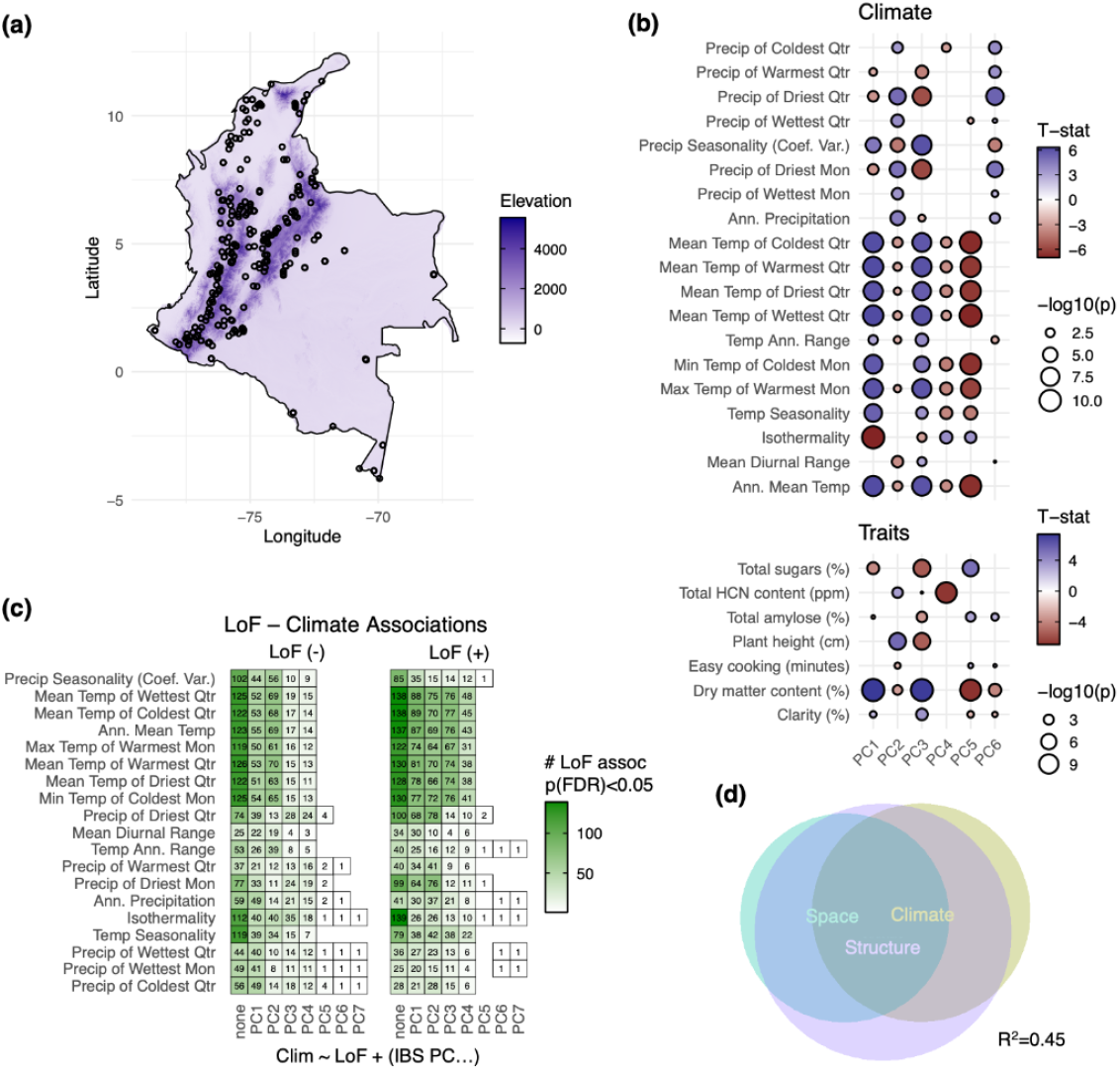
Collinearity of Environmental Adaptation and Population Structure in Colombian Cassava Landraces. **(a)** Geographic locations of Colombian landrace samples analyzed in this study. **(b)** Correlation of climatic variables and agronomic traits with the top six genetic principal components (PCs). Circle size indicates the significance level of the correlation, while color represents the direction (positive or negative). **(c)** The number of significant associations between LoF variants and climate variables decreases as more PCs are included to control for population structure. Numbers in each box indicate the count of significant associations after false discovery rate (FDR) correction. **(d)** Partitioning of genetic variance through redundancy analysis (RDA), quantifying the contributions of population structure, environmental variation (climate), and geographic location (space). Together, these factors explain 45% of the observed genetic variation.

Linkage disequilibrium (LD) decay rates vary among different populations and reflect their different demographic history. Overall, the linkage disequilibrium in cassava has a fast decay in 20 kb. African breeding lines decay to an average R^2^ of ∼0.21 in 20 kb, whereas South American accessions decay to an average R^2^ below 0.2 in 20 kb. Although differences in sequencing methods and depths among previous studies may introduce bias, we observed that LD in Colombian landraces is lower and decays more rapidly than in African and South American breeding lines (**Fig. 1c**), indicating higher genetic diversity and/or more abundant historical recombination events in the landraces. As expected, LD is lowest in the wild relatives, consistent with their high genetic diversity. Genetic assignment analysis using ADMIXTURE reveals a more complex genetic structure in Colombian landraces. At K = 5, these landraces comprise five distinct ancestral components, each predominant in different subsets of accessions. In contrast, African breeding lines are largely dominated by a single ancestral group, which again implies a strong bottleneck effect and their potential demographic origin (**Fig. 1d**).

The identity-by-state (IBS) matrix derived from the VCF reveals that both Asian and African breeding lines have a higher genetic similarity to lowland (<500 Meters Above Sea Level, MASL) South American accessions than to highland (>500 MASL) accessions (**Fig. S2a**). These observations suggest that the African and Asian breeding accessions likely originated from genotypes with closer genetic relatedness to modern landraces found in the lowland regions of South America. Pre-adaptation to higher temperatures may have contributed to the success of these accessions in the African breeding program. The IBS analysis between domesticated cassava and other species or subspecies indicates a close relationship with *flabellifolia, peruviana*, and *tristis* (**Fig. S2b**), consistent with findings from the previous phylogenetic study (Simon *et al*., 2022).

Interestingly, a small subset of Colombian landraces with the lowest PC1 values cluster tightly together and are clearly separated from the rest of the samples in the principal component space (**Fig. 1b**). These accessions correspond to individuals that are overwhelmingly composed of a single ADMIXTURE group and show little evidence of genetic mixing with other populations (**Fig. 1d**). Their pairwise IBS scores are exceptionally high, ranging from 0.9977 to 0.9992, indicating that they are nearly genetically identical. Notably, these accessions are not technical replicates, but rather distinct landrace accessions collected from different geographic regions (**Fig. S3**). This high degree of genetic similarity across geographically diverse samples could reflect the spread of a clonally propagated or strongly bottlenecked lineage, potentially due to human-mediated dispersal.

Additionally, we carried out an extensive analysis of genetic diversity and differentiation among populations. The wild relatives population displays the highest genetic diversity, with an average genome-wide π_WR_ of 1.90 × 10^−2^. The genetic diversity of South American cassava (π_SA_ = 8.91 × 10^−3^), which mostly consists of Colombian landraces, is higher compared to the African cassava population (π_A_ = 8.22 × 10^−3^), mostly comprising local breeding lines. This result further implies bottleneck effects throughout cassava domestication history. We also observed several genomic windows where π_WR_/π_SA_ or π_WR_/π_A_ is significantly higher, and those regions are likely subject to selection during domestication (**Fig. S4**). Similar π_landrace_/π_improved_ or π_wild_/π_landrace_ peaks have been observed in maize membrane trafficking genes that were selected during maize domestication and improvement (Zheng *et al*., 2022). The genome-wide average F_ST_ between South American cassava and African cassava is 0.0471, indicating relatively low genetic differentiation between these two cultivated or pre-breeding populations. In contrast, the F_ST_ between South American cassava and wild relatives is higher at 0.134, suggesting a greater genetic divergence. Similarly, the F_ST_ between African cassava and wild relatives is 0.125, also reflecting a significant degree of genetic differentiation.

### Environmental Adaptation of Colombian Landraces

While the large collection of cassava landraces represents vast untapped genetic diversity, they are also a key resource representing environmental adaptation. These landrace cassava clones originate from locations across vast environments in Colombia (**Fig. 2a**). Notably, the field sites from which these individuals originate correspond to varied climates. The largest impact of the climatic variation among individuals is driven by elevation distribution around the Andes mountains running through Colombia (**Fig. 2a, Fig. S1**). The lowland regions experience higher temperatures and lower precipitation than the highland areas.

The environments in which these clones are cultivated likely reflect adaptive selection histories, as these landraces have been grown in local conditions for hundreds of years. We observed strong correlations between environmental variables and the top genetic principal components. Notably, PC1 and PC3 were positively correlated with temperature, while PC2 showed a positive correlation with precipitation (**Fig. 2b**). Redundancy analysis (RDA), which assesses the proportion of genotypic variation explained by environmental, spatial, and population structure variables, revealed that these factors together account for a substantial portion of genetic variation (R^2^ = 0.45). While the contribution of population structure is expected, partial RDA indicates that climate alone explains a significant fraction of this variance (**Fig. 2d**). The shared variance among climate, space, and population structure underscores the challenge of disentangling the genetic basis of climate adaptation, and can be particularly problematic for GWAS. We further tested for associations between loss-of-function alleles and climate variables, finding that the number of significant associations declined as more genetic PCs were included in the model. When the top seven PCs were controlled for, few associations remained (**Fig. 2c**). Nonetheless, these results suggest that strong genetic correlations with environmental variables persist.

### Gene Loss-of-Function in Cassava

In an effort to enrich for impactful genetic elements that may explain climate adaptation, we evaluated the distribution of gene loss-of-function alleles across the cassava landraces. LoF mutations represent a subset of genetic variants that are expected to have large functional effects on gene activity. In total, we found over 11k variants segregating across the landraces categorized as causing a large disruptive effect by SnpEff (Cingolani *et al*., 2012). We used protein structure to reduce false positive rate of classifying LoF, by disregarding mutations that are less likely to impact structured (folded regions with low available surface area), functional regions of the protein (**Fig. 3a, d&e**). The relative importance of peripheral, disordered regions of proteins is a subject of current scientific inquiry, but we are choosing to disregard them for this analysis. The LoF mutations are confined to ∼6.5k genes (∼20% of the proteome) with many genes containing multiple segregating variants (**Fig. 3b**). We collapsed the LoF classification across all genes to a singular functional classification for each gene (**Fig. 3d&e**) and evaluated the frequency of LoF (**Fig. 3c**). After filtering out rare, low frequency functional variation (MAF > 5%), over 2k genes remained.

**Figure 3.**
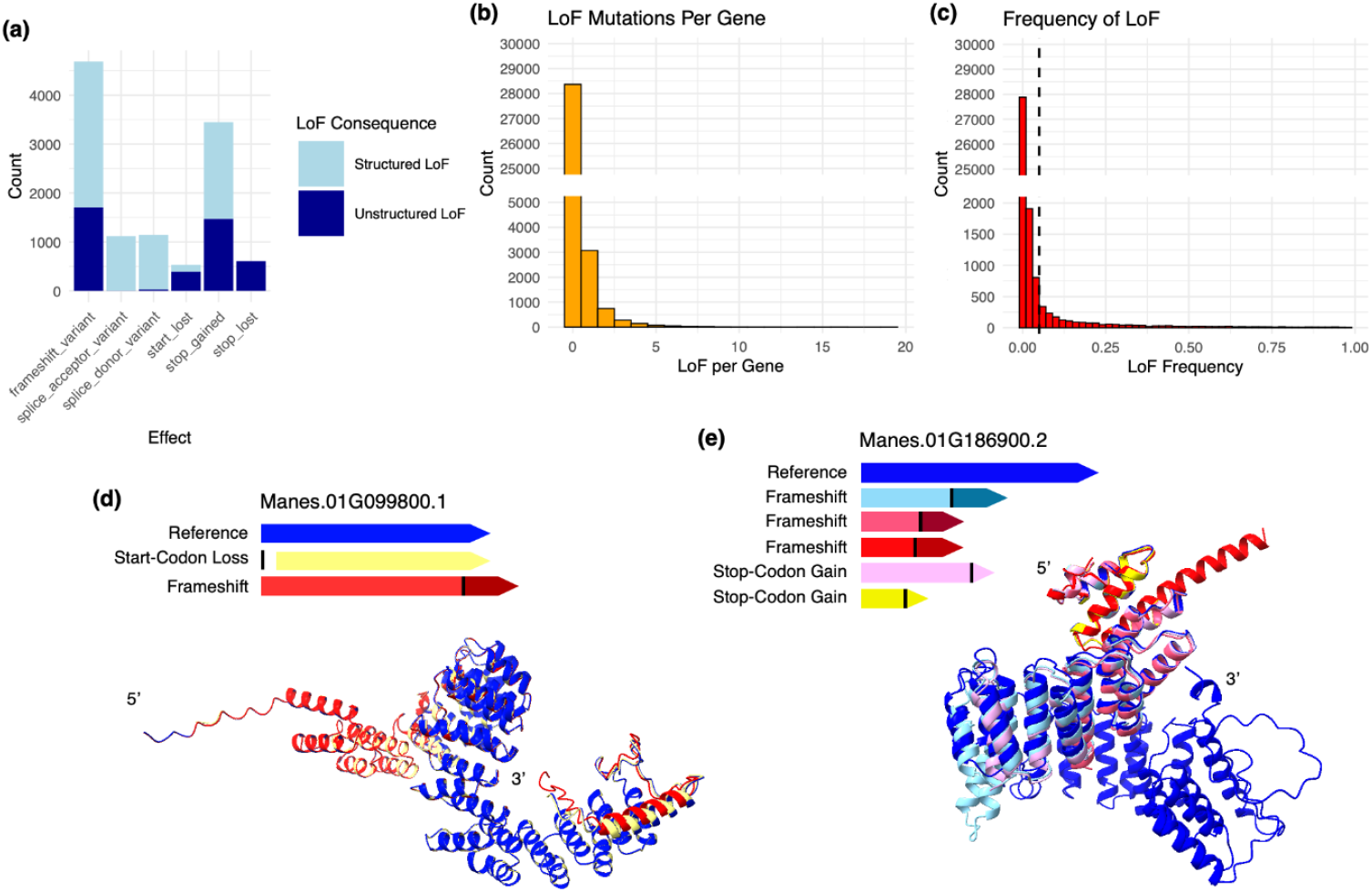
Types, Distribution, and Structural Impact of Gene Loss-of-Function Variants in Cassava. **(a)** Histogram of gene LoF variant types predicted by SnpEff in this study. Frameshift mutations and premature stop codons account for the majority of predicted LoF events. The functional impact of LoF variants is further assessed through protein structure predictions. “Structured LoF” refers to variants likely occurring within structured protein regions, thus more likely representing true LoF events. **(b)** Histogram showing the number of independent LoF variants identified per gene. **(c)** Histogram of LoF variant frequencies per gene across the studied population. The dashed line marks the 5% minor allele frequency (MAF) threshold commonly applied in association studies. **(d & e)** Illustrative diagrams of LoF variants for two example genes, comparing their their predicted protein structures relative to the reference gene model. Black bars indicate the locations of LoF mutations within the gene body. Protein structure models depict the impact of LoF variants, highlighting deviations from the reference structures.

Cassava is a highly heterozygous crop that experiences severe inbreeding depression (Ramu *et al*., 2017; Long *et al*., 2025), making the accumulation of homozygous deleterious mutations particularly detrimental. To investigate the relationship between inbreeding and deleterious variation, we calculated the inbreeding coefficient (F) for each individual in our dataset using genome-wide genotype data. We observed a significant negative correlation between the number of LoF alleles per individual and their inbreeding coefficient (Spearman’s ρ = –0.46, **Fig. S5a-c**). This suggests that inbred individuals tend to carry fewer LoF alleles, consistent with purging of deleterious mutations through inbreeding.

To investigate the functional role of LoF variants in evolution and adaptation of cassava, we performed gene ontology (GO) analysis on genes retaining LoF alleles. Our results reveal that LoF-tolerant genes in cassava are significantly enriched for immune response functions (**Fig. 4**), consistent with previous findings in *Arabidopsis* (Zhao *et al*., 2025). These genes include compelling candidates that may be leveraged to engineer disease resistance in cassava through genetic manipulation. For example, Manes.16G055900 is homologous to Arabidopsis AT2G35110 (*HEM1*), a global translational regulator of plant immunity. Loss of *HEM1* leads to exaggerated cell death that restricts bacterial growth and enhances immunity (Zhou *et al*., 2023). Manes.08G172100 shares homology with AT1G28380 (*NSL1*) and AT1G29690 (*NSL2*), which participate in the salicylic acid (SA)-mediated programmed cell death pathway in plant immunity (Morita-Yamamuro *et al*., 2005; Noutoshi *et al*., 2006; Murakoshi *et al*., 2024). Mutants with *NSL1* knockout constitutively activate defense responses (Noutoshi *et al*., 2006). Manes.03G015900, homologous to AT1G51940 (*LYK3*), is involved in suppressing immune responses in the absence of pathogens or following abscisic acid treatment. *LYK3* loss-of-function mutants exhibit increased resistance to *Botrytis cinerea* and *Pectobacterium carotovorum* (Paparella *et al*., 2014). Manes.08G125011, related to AT5G61900 (*BON1*) and AT1G08860 (*BON3*) in *Arabidopsis*, may also function in plant immunity, as *BON1* and *BON3* negatively regulate several resistance (R)-like genes (Li *et al*., 2009), and their knockout in both *Arabidopsis* and rice has been shown to enhance resistance to bacterial and fungal pathogens (Li *et al*., 2009; Yang *et al*., 2017).

**Figure 4.**
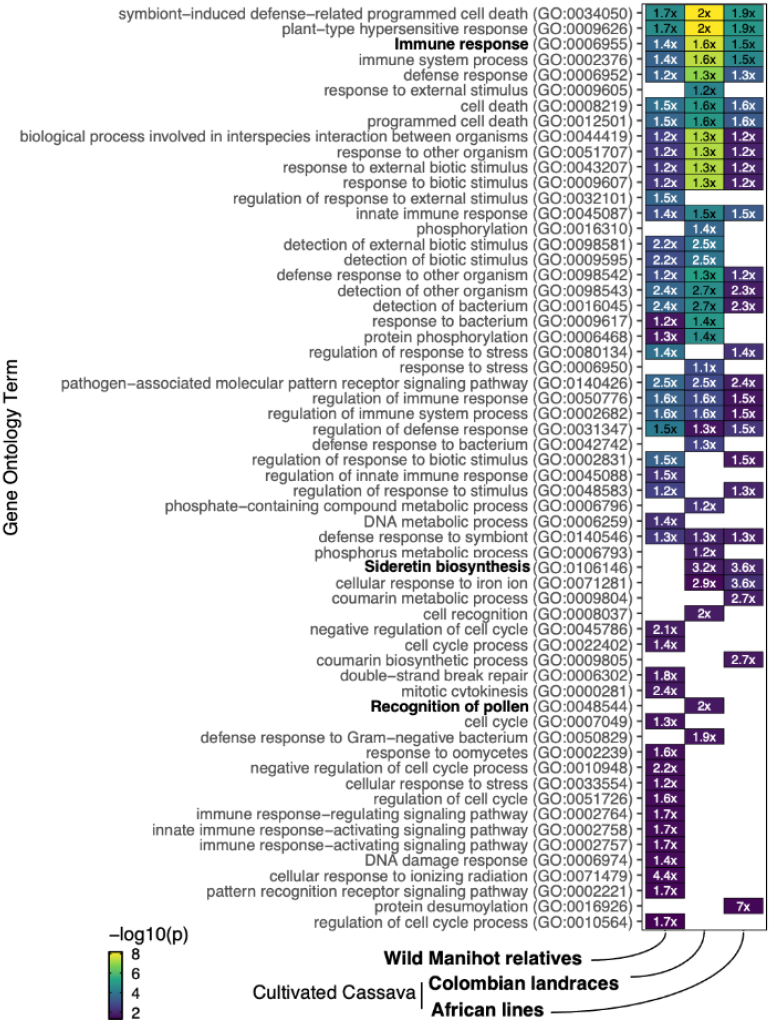
Gene Ontology (GO) Enrichment Analysis of LoF-Tolerant Genes in Cultivated Cassava and Wild Relatives. Significantly enriched GO terms for LoF-tolerant genes are displayed, ranked by significance. The numbers within each box represent the fold enrichment of the corresponding GO term. Notable biological processes are highlighted in bold for emphasis.

In addition to the enrichment for immune response, we observed a 3.6-fold enrichment for sideretin biosynthesis and a 2.7-fold enrichment for the coumarin metabolic process in African breeding lines, respectively (**Fig. 4**). Coumarins are a class of secondary metabolites that play a central role in iron (Fe) solubilization and uptake, particularly under alkaline soil conditions (Robe *et al*., 2021). Among them, compounds such as fraxetin and sideretin have been shown to facilitate Fe acquisition, with their efficacy varying depending on soil pH (Rajniak *et al*., 2018; Paffrath *et al*., 2024). Beyond their physiological functions, coumarins are also agriculturally significant. Scopoletin, the chemical precursor to sideretin, and its glucoside, scopolin, have both been implicated in post-harvest physiological deterioration (PPD) in cassava (Buschmann, 2000; Bayoumi *et al*., 2008, 2010). Specifically, scopoletin oxidation leads to tissue discoloration, rendering roots unpalatable and reducing their marketability (Liu *et al*., 2017). We found an enrichment of LoF alleles in genes associated with sideretin biosynthesis, including homologs of *CYP82C2, CYP82C3*, and *CYP82C4* from *Arabidopsis thaliana*. Notably, CYP82C4 catalyzes the hydroxylation of fraxetin to produce sideretin. Additionally, Manes.07G034800 was identified as a homolog of AT1G55290 (*F6’H2*), AT3G13610 (*F6’H1*), and AT3G12900 (*S8H*). Of these, *F6’H1* is involved in the biosynthesis of scopoletin from feruloyl-CoA, while *S8H* hydroxylates scopoletin to form fraxetin (Rajniak *et al*., 2018). Whether the accumulation of LoF alleles in coumarin-related genes reflects selection against undesirable post-harvest discoloration or adaptation to varying soil conditions remains unresolved and motivates further investigation.

Moreover, we found that LoF-tolerant genes are enriched for functions related to pollen recognition (**Fig. 4**). This result is consistent with the previous report suggesting that domestication and the shift to clonal propagation in cassava have relaxed selection on sexual reproduction, allowing mutations to accumulate in pollen-associated genes (Long *et al*., 2025). Interestingly, although there is a genome-wide negative correlation between the number of LoF alleles per individual and their inbreeding coefficient, this correlation is markedly less negative or even positive for genes within significantly enriched GO categories (**Fig. S5**).

The persistence of LoF alleles in these genes, despite inbreeding, suggests they may confer potential adaptive or beneficial effects.

### Genome-Environment Association

In order to pinpoint specific genetic variants responsible for climate adaptation, we performed genome-wide association between genetic variants of the landraces genotypes and many environmental variables derived from their geolocation. We used three different classifications of genetic variants for environmental association including all variants, variants predicted to result in gene LoF (**Data S1**), and the consolidated gene LoF burden (**Data S2**).

Many climatic variables were tested (**Table S2**), while we show the results to two primary dimensions of environmental variation: temperature and precipitation (**Fig. S6**). Genetic associations with annual temperature, an environmental variable of major interest with impending climate change, found a significant locus on chromosome 6 among all variants and LoF variants. This locus, however, was not retained when considering the LoF burden classification that incorporates predicted effects on protein structure. Associations with annual precipitation showed multiple significant variants across the cassava genome. A handful of these variants corresponded to LoF variants and the LoF burden of two specific genes when controlling for the impact on protein structure. These two genes, Manes.09G029200 and Manes.10G053700, show opposite effects in their correlation with high and low precipitation, respectively. In addition to climatic variables, we also tested a few phenotypes that were collected from historical data. We found a significant association with high sugar content corresponding to the LoF burden of a gene on chromosome 6. This gene is annotated as a glycosyltransferase, suggesting that its LoF may impact the ability to transport sugars outside of the roots. While high sugar content is generally not a valued trait for cassava production, it is an interesting proof of concept for the utility of LoF association analysis.

We also used redundancy analysis to look for genes responsible for climate adaptation. The RDA method uses collapsed variables space to assess which climate variables explain mutations in the genome, somewhat reversed from a typical genome-wide association analysis. When looking at what genes are most highly explained by climate variables, one gene, Manes.01G186900.2, proved significant (**Fig. S7**). This gene was among the most significantly associated genes with annual temperature, but did not pass any significance thresholds in the GWAS analysis. The LoF in this gene is also derived from multiple segregating mutations (**Fig. 3e**), making it a particularly compelling candidate for further investigation.

## Discussion

In this study, we characterized previously untapped genetic diversity in Colombian cassava landraces. Our analysis of population structure and genetic diversity aligns with the known domestication history of cassava, which originated in South America and was later introduced into Africa and Asia. Principal component analysis reveals that African and Asian breeding lines partially overlap with South American accessions, suggesting a genetic connection to the South American gene pool underlying modern cassava cultivars. The nucleotide diversity of African cassava is 8.22 × 10^−3^, which is higher than the previously reported nucleotide diversity (π = 3.6 × 10^−3^) by Ramu *et al*. (2017). This discrepancy may reflect methodological differences, as nucleotide diversity in the previous study was estimated using VCFtools, which may have limitations under certain conditions (Korunes & Samuk, 2021). Cassava is an outbreeding plant species, and despite being a clonally propagated crop, the nucleotide diversity of Colombian landraces (∼8.91 × 10^−3^) is higher than previously reported π in inbreeding crops such as soybean landraces (∼1.43 × 10^−3^) (Hyten *et al*., 2006), sorghum landraces (∼2.2 × 10^−3^) (Hamblin *et al*., 2006), wild Emmer wheat (∼2.7 × 10^−3^) and cultivated wheat (∼8 × 10^−4^) (Haudry *et al*., 2007), common wheat landraces (∼9.45 × 10^−4^) (Niu *et al*., 2023), rice wild relatives (∼6.4 × 10^−3^) (Zhu *et al*., 2007), comparable to the previously reported π in different maize landrace populations (∼6-10 × 10^−3^) (Tittes *et al*., 2023), which is also an outbreeding crop. The high nucleotide diversity in cassava landraces is a valuable asset that breeding programs can leverage to develop new varieties with desirable traits such as disease resistance, drought tolerance, and improved nutritional content.

By sampling genetic diversity of cassava landraces across the environmental landscapes of Colombia, we aimed to capture potential adaptation to their local environment. This hypothesis that the location in which these landraces were sampled may correspond to environmental adaptation relies upon the assumption that farmers retained those varieties that performed best and were well adapted to their respective regions (Janzen *et al*., 2022). The correlation of population structure derived from genetic relationships and the climatic variables represents the underlying double-edged sword of this research approach. In contrast to the findings by Kistler *et al*. (2025), which found minimal geographic population structure in cassava when examined at large spatial scales across the Americas attributed to clonal propagation and human-mediated dispersal, our data reveal pronounced population structure among Colombian landraces. We also found that population structure showed high correlation to many environmental variables, which is consistent with both adaptation and demographic history. Unfortunately, such structure makes it difficult to identify specific adaptive causative mutations. Controlling for population structure in our associations attempts to reduce false positive associations due to structure-environmental correlations, meanwhile, we acknowledge that this is also likely reducing our ability to capture true positive relationships (Lasky *et al*., 2015).

In addition to conventional genotype–environment association using SNP data, we developed a function-based GWAS approach that associates LoF variants and LoF burdens. By collapsing independent LoF alleles into a single allele state, we aim to overcome allelic heterogeneity, which has limited previous GWAS efforts. Moreover, traditional GWAS typically highlights genomic regions rather than pinpointing specific causative mutations, often leaving results difficult to translate into practical applications like gene editing. In contrast, our LoF association tests directly link phenotypes to gene function, enabling the identification of actionable gene-level targets. While population structure complicates the detection of causal variants, we propose that relaxing population structure controls can be justified to uncover candidate genes because of their testability via gene editing.

While LoF association tests offer valuable insights, some cautions are warranted. First, relying on a single reference genome may bias LoF detection, missing variation absent from the reference or structural variants (Zhou *et al*., 2022). Incorporating multiple references or pangenomes can mitigate this. Second, predicted LoF variants do not always cause phenotypic loss due to alternative transcription and/or splicing, compensatory mutations, or genetic redundancy (Singer-Berk *et al*., 2023). We applied protein structure prediction to reduce false positives, but factors like heterozygosity, dominance, and pseudogenes in cassava still complicate interpretation (Rausell *et al*., 2020). Lastly, LoF variants may be in linkage disequilibrium with causal variants, requiring careful analysis and functional validation to confirm causality.

Although our LoF analysis has yielded promising insights into the genetic basis of trait variation and climate adaptation in cassava, further work is needed to fully translate these findings into functional understanding and agronomic application. Experimental validation of candidate LoF genes, such as through CRISPR-based gene editing, will be essential to confirm their roles. Additionally, obtaining agricultural phenotypes in the field under diverse environmental conditions can help bridge the gap between genotype and environment, advancing our ability to harness LoF variation for crop improvement.

## Acknowledgments

We thank members of Monroe Lab for valuable discussions of this work. This work was supported by FFAR grant ICRC20-0000000014. Research was conducted at the University of California Davis, which is located on land that was the home of the Patwin people for thousands of years.

## Data availability

Figures, supplemental data, and code for this research are located at: https://github.com/KehanZhao/ColombianCassavaLandraces.

All raw genomic sequencing data generated in this study are publicly available through the NCBI Sequence Read Archive (SRA) under Bioproject accession number PRJNA1228154. Sample metadata for both newly generated and previously published data are provided in Table S1. For access to additional post-processed data formats, please contact the corresponding authors.

## Supplemental tables and data

Table S1. Meta information of 1,152 accessions included in this study

Table S2. Summary table of GWAS and RDA

Data S1. Predicted LoF variants in Variant Call Format

Data S2. Gene-level LoF burdens in Variant Call Format

## Supplemental figures

**Figure S1.**
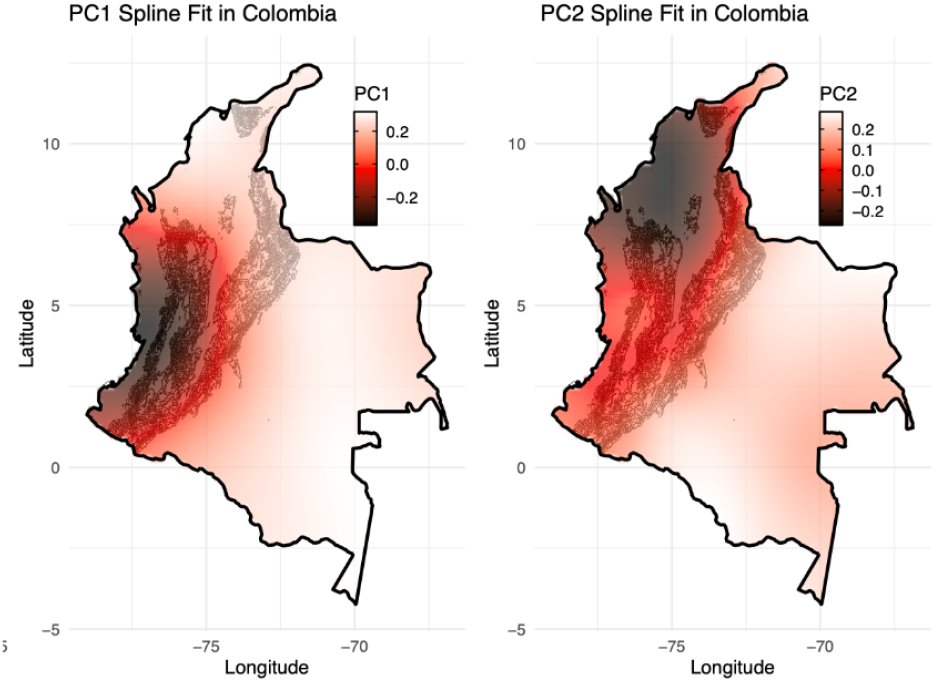
Correlation Between Principal Components and Geography. Map illustrating the geographic distribution of the first two genetic principal components (PCs). Lighter colors represent positive PC values, while darker colors indicate negative values. Notably, lower PC1 values are associated with the highland regions of the Colombian Andes.

**Figure S2.**
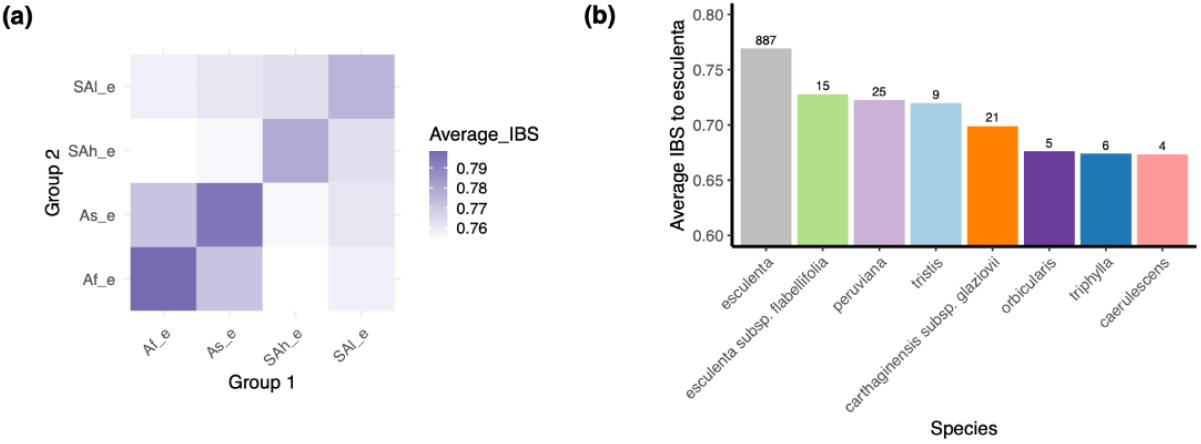
Identity-By-State (IBS) Relationship between Cassava Accessions and Close Relatives. **(a)** Average IBS between African *M. esculenta* (Af_e), Asian *M. esculenta* (As_e), South American highland *M. esculenta* (SAh_e), and South American lowland *M. esculenta* (SAI_e). (b) Average IBS of close relative species compared to *M. esculenta*. Numbers above bars indicate sample sizes.

**Figure S3.**
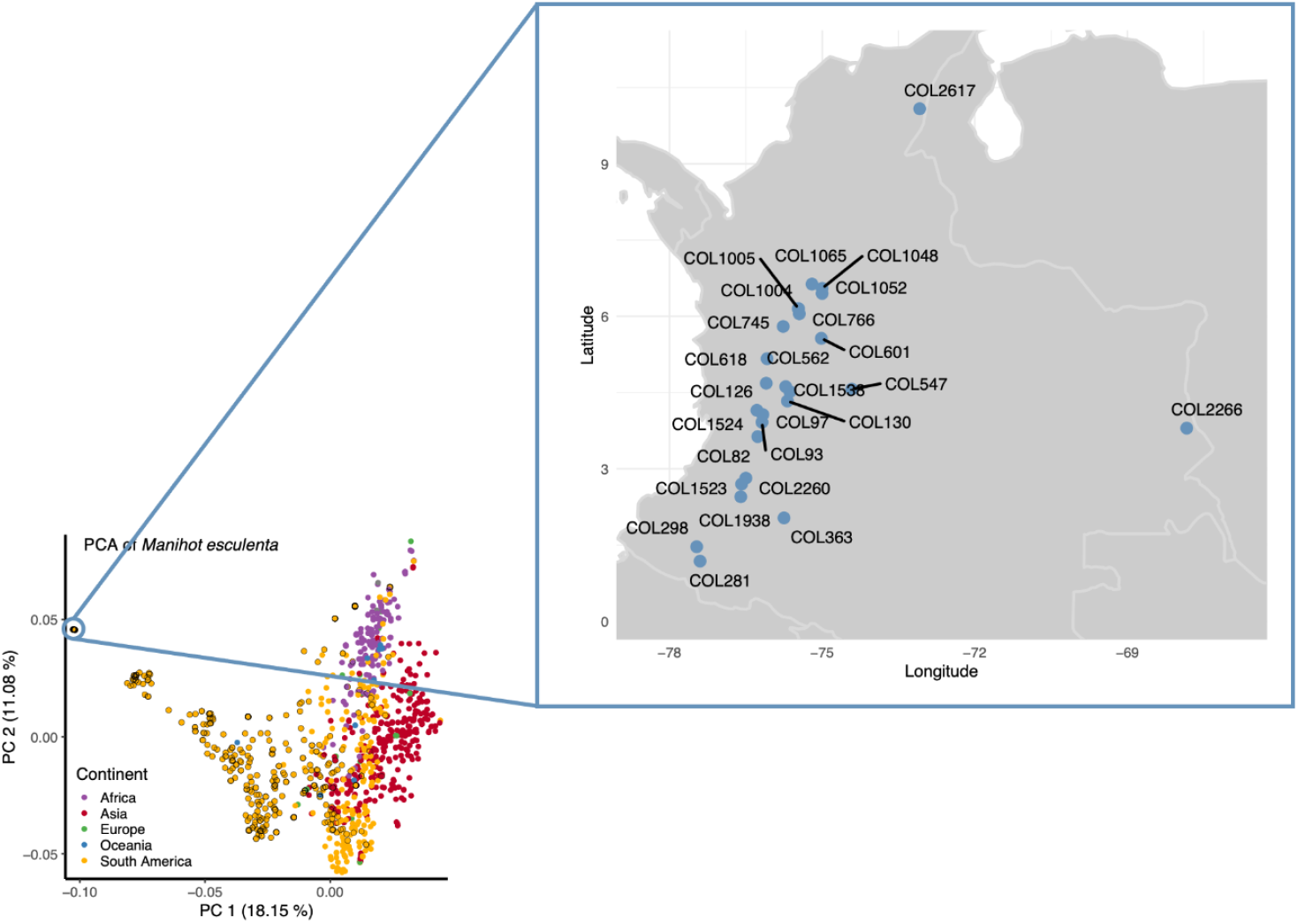
Geographic Location of Accessions with High Genetic Identity Similarity. Distinct landrace accessions collected from different geographic regions cluster tightly on principal component space. Those accessions have high pairwise IBS scores.

**Figure S4.**
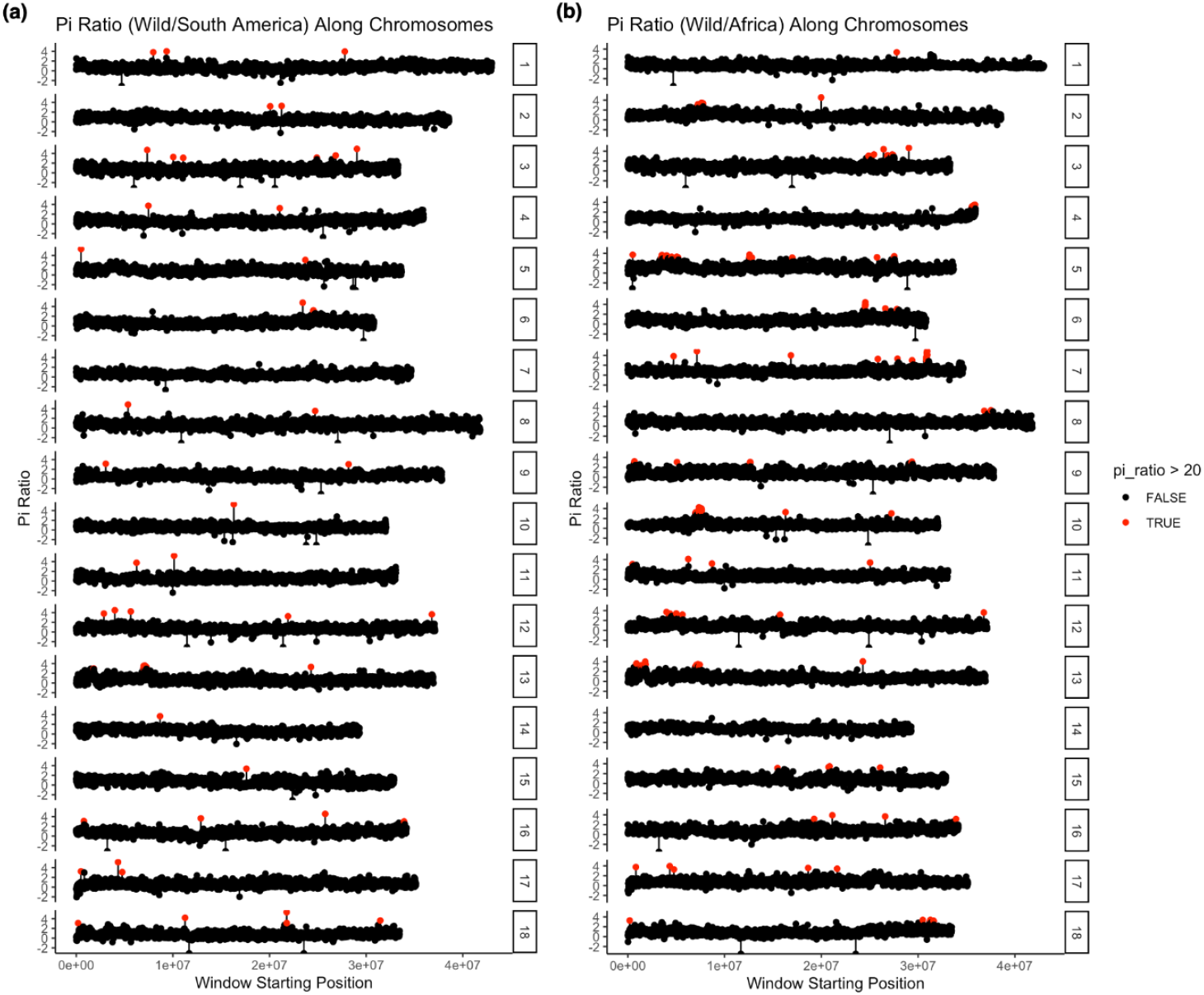
Nucleotide Diversity Ratio of Wild Relatives to Cultivated Lines. (a) Chromosome-wide log-transformed ratio of nucleotide diversity in wild relatives vs. South American lines (π_WR_/π_SA_ (b) Chromosome-wide log-transformed ratio of nucleotide diversity in wild relatives vs. African lines (π_WR_/π_A_). Red dots indicate genomic regions with elevated diversity ratios.

**Figure S5.**
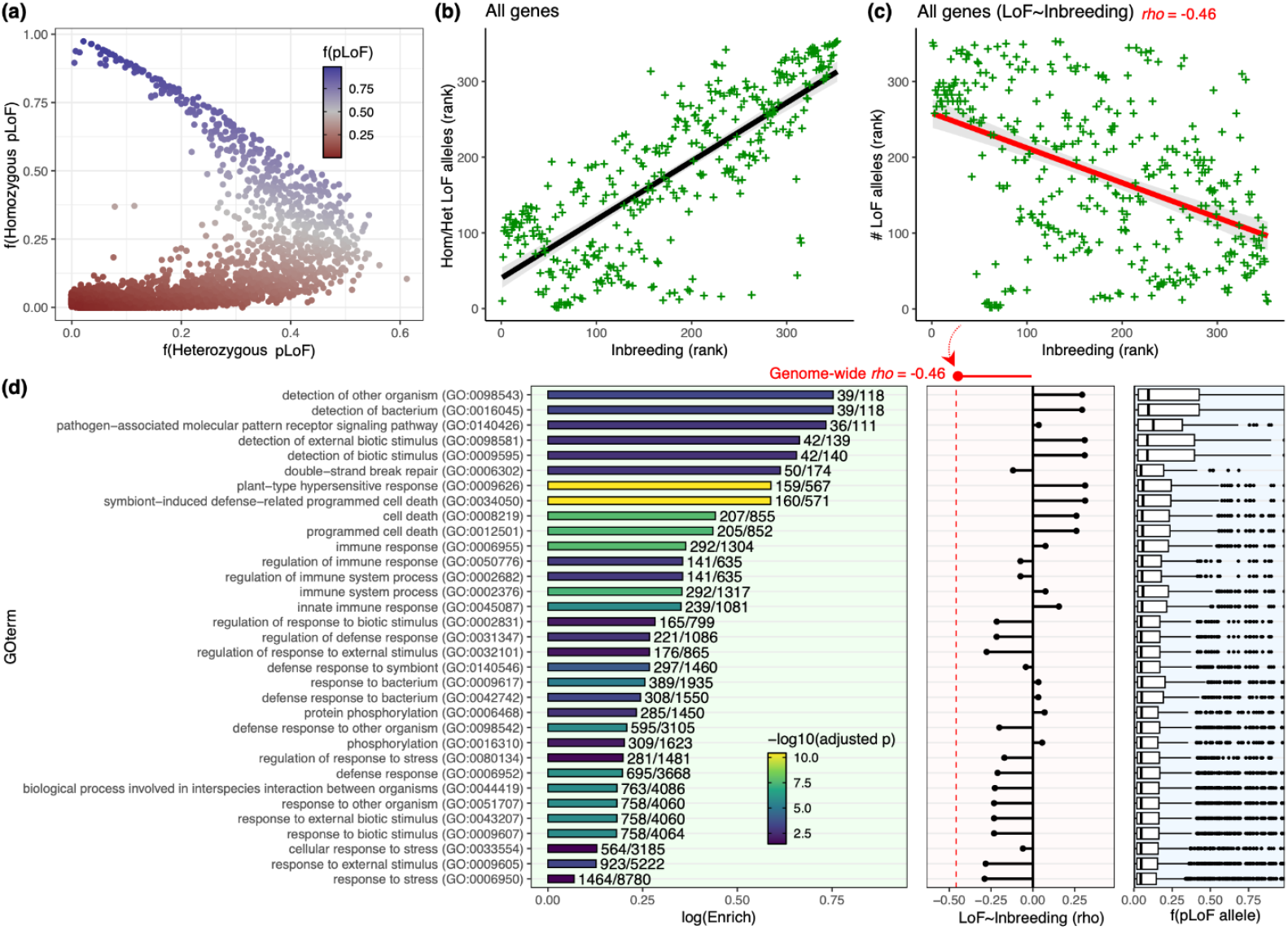
Relationship between Loss-of-Function (LoF) Variation and Inbreeding across the Genome. **(a)** Proportion of homozygous vs. heterozygous predicted LoF (pLoF) alleles per line. Each point represents an individual; points are colored by the total frequency of pLoF alleles in that line. **(b)** Positive correlation between inbreeding coefficient and the ratio of homozygous vs heterozygous pLoF alleles across accessions. **(c)** Negative correlation between inbreeding coefficient and total number of pLoF alleles per line (Spearman’s ρ = −0.46, shown in red). **(d)** Gene Ontology (GO) enrichment analysis of LoF-tolerant genes in cassava. Bar plot shows enriched GO terms and corresponding −log_10_(adjusted p-value), enrichment (log scale), and number of affected genes (numerator: number of LoF genes in term; denominator: total number of genes in term). Middle panel shows the correlation (p) between LoF allele count and inbreeding coefficient for each GO term; right panel shows the distribution of LoF allele frequencies in each GO term.

**Figure S6.**
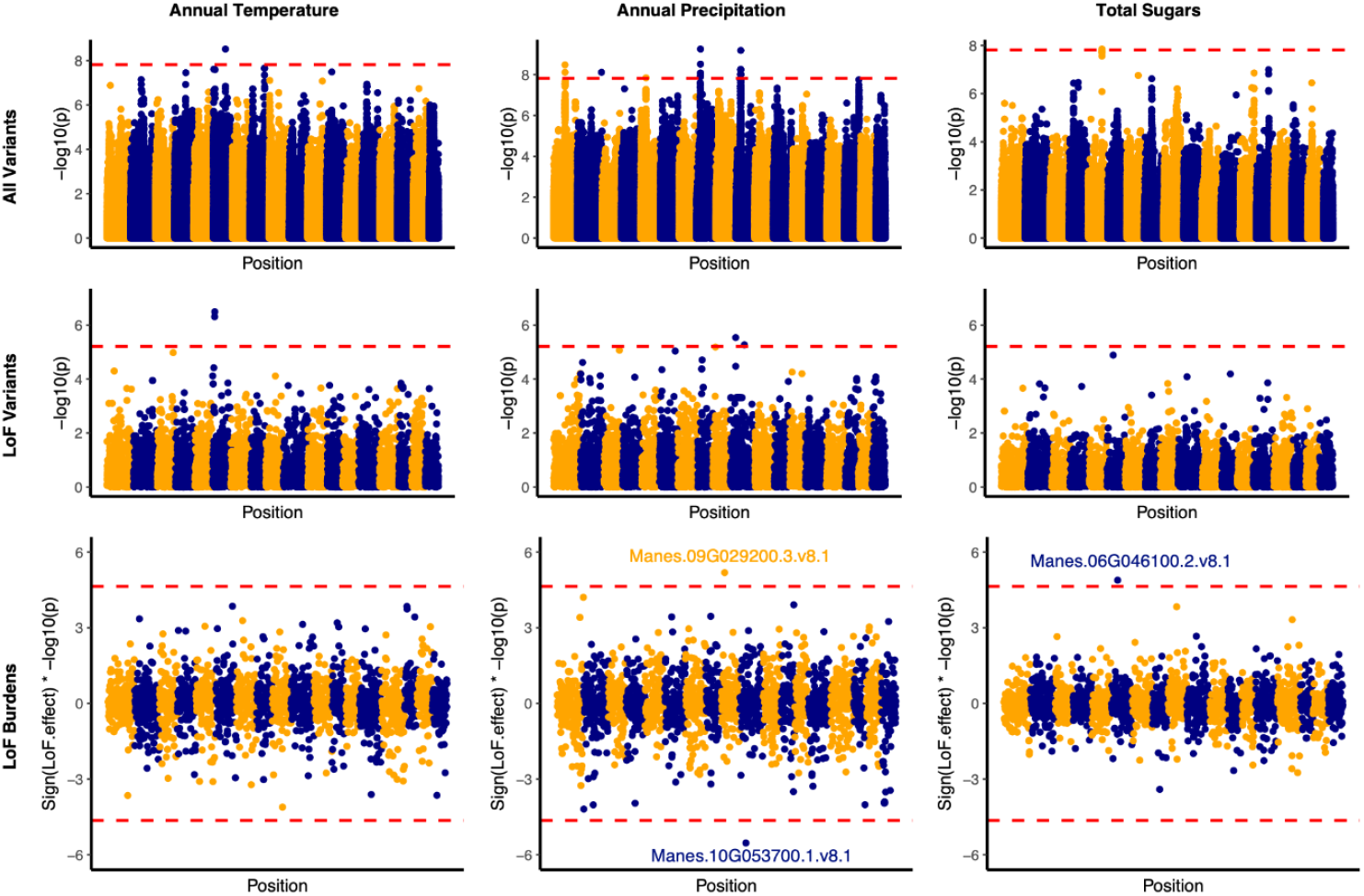
Manhattan Plots of Genome-Wide Association Tests Using All Variants. LoF Variants and LoF Burdens. Association results are shown for two key climate variables (annual temperature and annual precipitation) and for total sugar content. Points are colored alternately in yellow and blue to distinguish chromosomes. The red dashed line indicates the Bonferroni-corrected significance threshold. For LoF burden tests: the direction of the LoF effect is visualized by plotting −log_10_(p) values as positive or negative: points above zero indicate that gene LoF is positively associated with the climate variable or trait: points below zero indicate a negative association; significant associations surpassing Bonferroni threshold are labeled.

**Figure S7.**
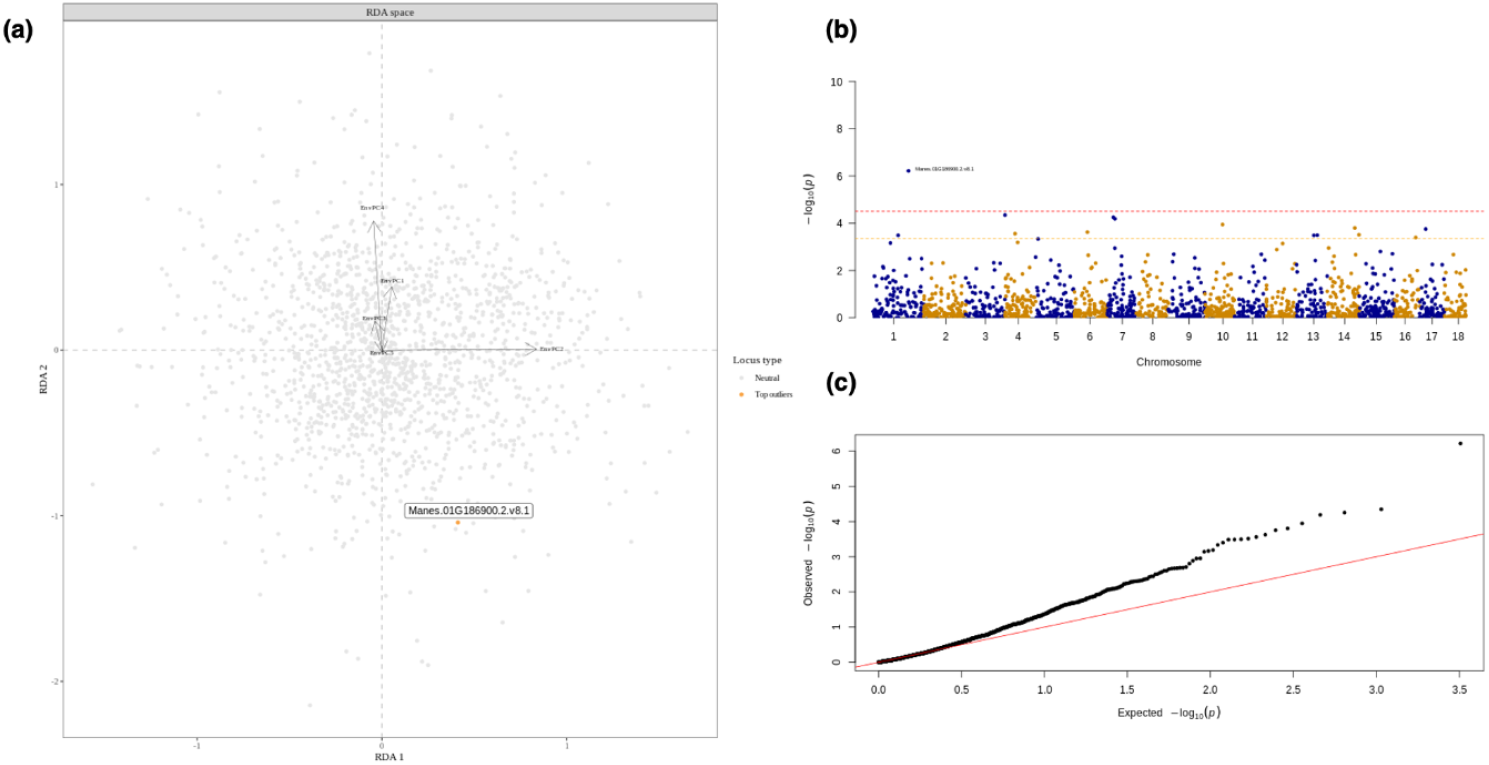
Genotype-Environment Association using Redundancy Analysis (RDA). **(a)** RDA biplot of Colombian cassava landraces. Each point represents a genetic locus, positioned according to its loadings on the first two constrained RDA axes (RDA1 and RDA2), which capture the greatest genetic variation explained by environmental predictors. Arrows indicate the direction and strength of correlation between each environmental principal component (EnvPC1−5) and genetic variation. Grey points represent putatively neutral loci, while the top candidate outlier locus (Manes.01 G186900.2) is highlighted in orange. **(b)** Manhattan plot showing the distribution of −log_10_-transformed p-values derived from Mahalanobis distances of each locus from the RDA centroid. Dots above Bonferroni-corrected p-value threshold (red dash line) or FDR-corrected p-value threshold (yellow dash line) indicate loci with significant associations to environmental gradients. **(c)** Quantile-quantile (QQ) plot comparing observed p-values from the Mahalanobis distance test to the expected uniform distribution under the null hypothesis. Deviation from the diagonal indicates loci with stranger-than-expected associations, supporting their candidacy as environmentally responsive outliers.

